# Apoptotic caspases silence spontaneous innate immune signals by specifically cleaving activated mitochondrial antiviral signalling protein (MAVS)

**DOI:** 10.1101/2025.02.25.640042

**Authors:** Sylwia Gradzka-Boberda, Ishita Parui, Pallab Chakraborty, Danielle Paige Anthony, Bhagya K. Puthussery, Arnim Weber, Dominik Brokatzky, Piero Giansanti, Julia Mergner, Rupert Öllinger, Roland Rad, Susanne Kirschnek, Ian E. Gentle, Georg Häcker

**Author notes:** These authors contributed equally: Sylwia Gradzka-Boberda, Ishita Parui, Pallab Chakraborty.

## Abstract

Caspases-9, -3 and -7 are activated in the mitochondrial apoptosis pathway and lead to the apoptotic phenotype. Caspases also function to limit inflammation upon apoptotic mitochondrial permeabilization through degradation of the signalling proteins cGAS, MAVS and IRF3. Cells and mice lacking caspases have higher interferon levels and are resistant to viral infection. We report that in unstimulated, non-apoptotic cells caspase-3 functions to cleave specifically activated MAVS and very likely cGAS. In unstimulated HeLa cells, constitutive caspase-9- and -3-but not 7-dependent proteolytic events were observed. Inhibition of the mitochondrial apoptosis pathway in various healthy cells induced type I interferon (IFN I) through increased cGAS activity in the absence of changes to cGAS levels. We observed enhanced MAVS-dependent signals upon RIG-I-like helicase stimulation in the absence of BAX, caspase-9 or caspase-3 or upon caspase-inhibition. During activation, MAVS forms complexes, and blockade of mitochondrial apoptosis signalling increased complex abundance in unstimulated and stimulated cells. MAVS complexes were more sensitive to caspase-degradation than the monomer, and mutation of caspase-3-cleavage sites in MAVS spontaneously increased complex formation. Inhibition of voltage-dependent anion channel 1 (VDAC1) oligomerization blocked BAX/BAK- and caspase-regulated IFN induction, suggesting a stimulating role of leakage of mitochondrial DNA. We propose that low level, spontaneous activity of the mitochondrial apoptosis pathway, through specific caspase-3-mediated cleavage of only active signaling proteins, counteracts mitochondrial release of nucleic acids to reduce inflammation in the absence of infection. Caspase-3 therefore has a novel function in conformation- and activation-specific cleavage of substrates.

## Introduction

Cell death by apoptosis is often the result of mitochondrial permeabilization: the mitochondrial intermembrane space proteins cytochrome *c* and SMAC are released into the cytosol and activate caspases, which cleave numerous cellular proteins and are involved in the disposal of the dying cell (*1, 2*). More recently, it has become clear that mitochondria contain various molecules that can, when released, be recognized by cytosolic receptors and drive an inflammatory response. The best characterized molecule here is mitochondrial DNA (mtDNA) (*3*), which alongside mt (double-stranded) RNA (*4*) can be recognized by cytosolic pattern recognition receptors. Additional molecules of mitochondrial origin may be recognized by the same cell or, upon cell death, by neighbouring cells (*5, 6*). Apoptosis is, however, unlike other forms of cell death, non-inflammatory. This makes a mechanism necessary that blocks the pro-inflammatory signals that are generated by the release of mitochondrial contents. Intriguingly, caspase-mediated cleavage of signalling molecules has been identified as this anti-inflammatory mechanism. Inhibition or deletion of cell death caspases makes apoptosis inflammatory: a cell undergoing apoptotic mitochondrial permeabilization while prevented to activate caspases produces type I interferon (IFN I) (*7*) and NF-κB-dependent cytokines and chemokines (*8*). Even at steady state, caspase-9 (*9, 10*) and caspase-3 (*10*) deficient cells show enhanced production of IFN I and enhanced resistance to viral infection.

This suggests that cell death caspases cleave essential signalling proteins of innate immunity. Mitochondrial release of cytochrome *c*/SMAC activates caspase-9, which in turn activates the effector caspases 3 and 7. Caspase-3 and -7 have almost identical peptide substrate preferences (*11*), and they have overlapping but not completely identical protein substrates in cells (*2*), and potentially differing functions (*12*). To counter inflammatory signalling, caspase-3 cleaves the key signalling proteins cGAS, MAVS and IRF3 in human cells, with a role for caspase-7 only in mouse cells (*10*). During apoptosis induced by specific stimuli or by viral infection, these proteins were substantially degraded, presumably making them unavailable for signalling (*10*). This can explain the non-inflammatory nature of cell death. Because cells lacking caspase-activity will still die upon mitochondrial permeabilization, it has been proposed that the relevant function of cell death caspases is not, or at least not exclusively, the dismantling of the dying cell but the inhibition of inflammatory signalling (*13*).

These data make a cogent argument how caspases prevent inflammation during apoptosis. They do not however explain the higher spontaneous IFN I production in caspase-deficient cells (*9*). The finding suggests that caspases are spontaneously active in healthy cells and spontaneously degrade the signalling proteins, but caspase-generated fragments of cGAS or MAVS have not been found in non-apoptotic cells (*10*). Spontaneous non-lethal caspase activity, on the other hand, appears to exist. Cells can tolerate considerable amounts of caspase activity (*14*). Many reports have suggested physiological non-lethal roles of caspase-3, particularly in cell proliferation and tissue differentiation (*15, 16*). More recently, spontaneous, sub-lethal activity of the mitochondrial apoptosis pathway (*17*) has been identified in human cancer cell lines and appears to occur in mouse intestinal organoids (*18, 19*). In analogy to the situation during apoptosis, such spontaneous caspase activity might act to downregulate pro-inflammatory signalling, so caspase-deficient cells would have higher IFN I production. But how would this work? During apoptosis, cGAS/MAVS/IRF3 are degraded and presumably are simply not available for signalling but such degradation does not occur in non-apoptotic cells. Even though small levels of cleavage of the proteins at steady state might go unnoticed, there is no obvious explanation how such hard-to-detect potential cleavage of the proteins could reduce signalling intensity.

In this study, we investigated the mechanism of the down-regulation of pro-inflammatory signalling by low level, non-lethal activity of caspase-3. We found that the regulation of IFN I activity was due to the effect of caspase-3 on cGAS and to a smaller degree on MAVS, while ligand-driven activity of MAVS regulated IL-6-secretion and was controlled by caspase-3 in live cells. We identified mtDNA, spontaneously released through VDAC1, as a ligand driving cGAS activity. Strikingly, spontaneously active caspase-3 did not cleave the MAVS monomer distinguishably. Rather, it specifically reduced the amount of MAVS signalling complexes. Caspase-3 activity did not require a pro-apoptotic stimulus but was continuously generated by the mitochondrial apoptosis effector BAX. The results identify a function of spontaneously active caspase-3 in tuning the cellular reaction to infection. This activity appears to be required to silence the spontaneous inflammatory signals triggered by leakage of mitochondrial nucleic acids into the cytosol. More broadly, caspase-3 appears to have a function in conformation-specific substrate cleavage.

## Results

### In HeLa cells, caspase-9 and -3 but not -7 are active in steady state

We have previously reported evidence of spontaneous activity of the mitochondrial apoptosis pathway in human cell lines, starting from the identification of the activity of the caspase-activated DNase (CAD) in steady state (*18*). To identify the contribution of individual caspases, we measured cleavage activity towards the caspase-3/7 substrate DEVD-AMC in non-apoptotic cells from standard culture. Some cleavage activity was detectable (Fig. 1A); when investigating apoptosis, this would normally be considered ‘background activity’. Cells deficient in caspase-9 or caspase-3 harboured significantly less spontaneous DEVDase activity while there was only some inconsistent trend to less activity in caspase-7-deficient cells (Fig. 1A) (gene deficiency of all cells used in this study and not published before is documented in Fig. S1). We then identified cleaved caspase substrates in non-apoptotic cells using N-terminomics (i.e. identifying free N-termini of cellular proteins by MS-based proteomics), comparing control HeLa cells with HeLa cells individually deficient in caspase-3, -7 or -9. We identified a number of N-termini that are known to be generated by caspase-activity during apoptosis. Putative cleavage of a number of known caspase-3 substrates (such as vimentin, plectin, keratin-17) was observed in control cells but significantly less in cells deficient for either caspase-9 or caspase-3 (Fig. 1B; a novel substrate was also identified, Fig. S2A). This suggests that there is continuous cleavage of cellular proteins by caspase-3, which is continuously activated by caspase-9. No significant difference was seen in this assay in caspase-7-deficient cells, suggesting that caspase-7 is indeed not active at steady state. Spontaneous DEVD-cleavage activity was also seen in primary mouse macrophages differentiated from bone marrow in the presence of GM-CSF (Fig. 1C).

**Fig. 1.**
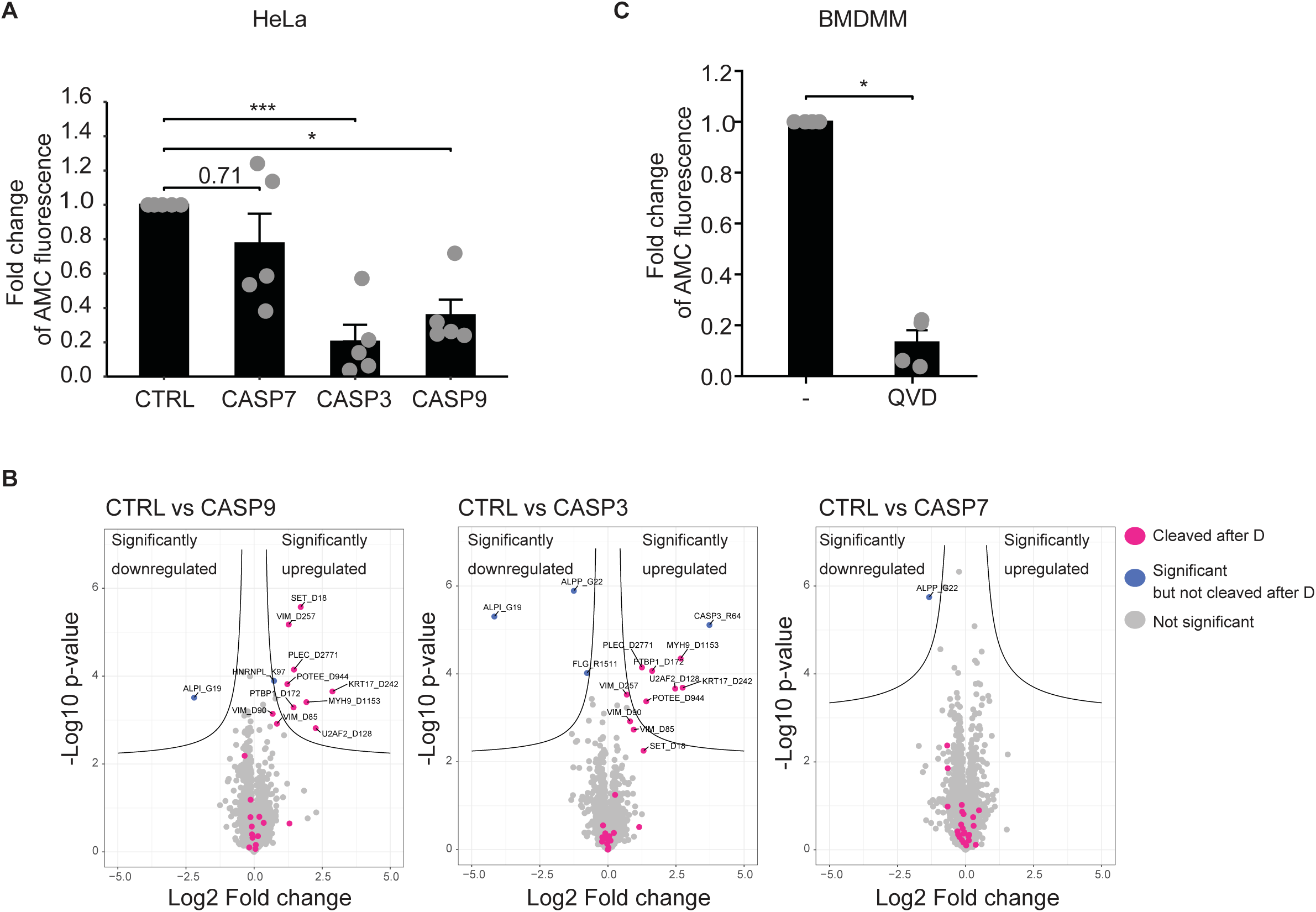
Constitutive activity of caspase-9 and -3 but not -7 in HeLa cells. **A** Cells from HeLa cell lines (CTRL, carrying a non-targeting gRNA or CRISPR/Cas9 mutants of the indicated genes) were lysed and analyzed for caspase-3-like activity using the Ac-DEVD-AMC substrate. Results from individual experiments are shown as dots and bars represent means of AMC fluorescence of five individual experiments. Error bars show the standard error of mean (SEM). Ns: p > 0.05, *, p < 0.05, ***, p < 0.001. The significance was tested by one-way ANOVA with Dunnett’s post hoc testing. **B** HeLa cell lines (CTRL, carrying a non-targeting gRNA or CRISPR/Cas9 mutants of the indicated genes) were evaluated by N-terminomics. Volcano plots show the distribution of protein N-termini identified in three separate experiments. N-termini generated by cleavage C-terminal to aspartic residues are highlighted in purple. The hyperbolic lines indicate the statistical significance threshold (adjusted p-values ≤ 0.05 with S0 = 0.1). **C** Mouse bone marrow derived monocytic macrophages (BMDMM) were differentiated with GM-CSF. Cells were treated with 10 µM QVD-OPh or DMSO (solvent control) for 24 h. Lysates were analyzed for caspase-3-like activity with the Ac-DEVD-AMC cleavage assay. Results from individual experiments are shown as dots and bars represent means of AMC fluorescence of four individual experiments (cells from four individual mice). Error bars show SEM. *, p < 0.05. The significance was tested by paired t-test.

### Enhanced cGAS activity by caspase deficiency but in the absence of changes to cGAS levels

It has been reported before that loss of caspase-9 or caspase-inhibition can cause the cGAS-dependent induction of IFN I in mouse embryonic fibroblasts (*9*). Confirming this concept, we identified an upregulated IFN response in caspase-9-deficient HeLa cells (Fig. 2A; Fig. S2B, C). Enhanced mRNA expression of IFN-β was seen in both caspase-9- and caspase-3-deficient cells (Fig. 2B), and similarly enhanced IFN-signalling was observed in BAX/BAK-, caspase-9 and caspase-3-deficient cells, measuring expression of the IFN response gene MX1 and phosphorylation of STING, TBK1 and IRF3 (Fig. 2C). Chemical caspase-inhibition had the same effect on the IFN response (Fig. 2G). Caspase inhibition activated the IFN system also in THP-1 human monocytic cells (Fig. 2D, Fig. S3A), as did the co-deletion of BAX and BAK in THP-1 cells (Fig. S3B, C). We also detected enhanced phosphorylation of TBK1 upon caspase inhibition in mouse bone marrow derived monocytic macrophages (Fig. 2E). Caspase-3-deficient HeLa cells showed enhanced cGAS activity (measured as the levels of the cGAS product cGAMP) (Fig. 2F). Deletion of cGAS abrogated the induction of IFN-β (Fig. S4A) and IFN signalling (Fig. 2G; Fig. S4B, C), and a similar effect was seen by deletion of STING (Fig. S4A, B, D). Inhibition of the mitochondrial apoptosis pathway in non-apoptotic cells therefore drives the activation of cGAS. This activation however occurs in the absence of any detectable changes in cGAS protein abundance (Fig. 2H).

**Fig. 2.**
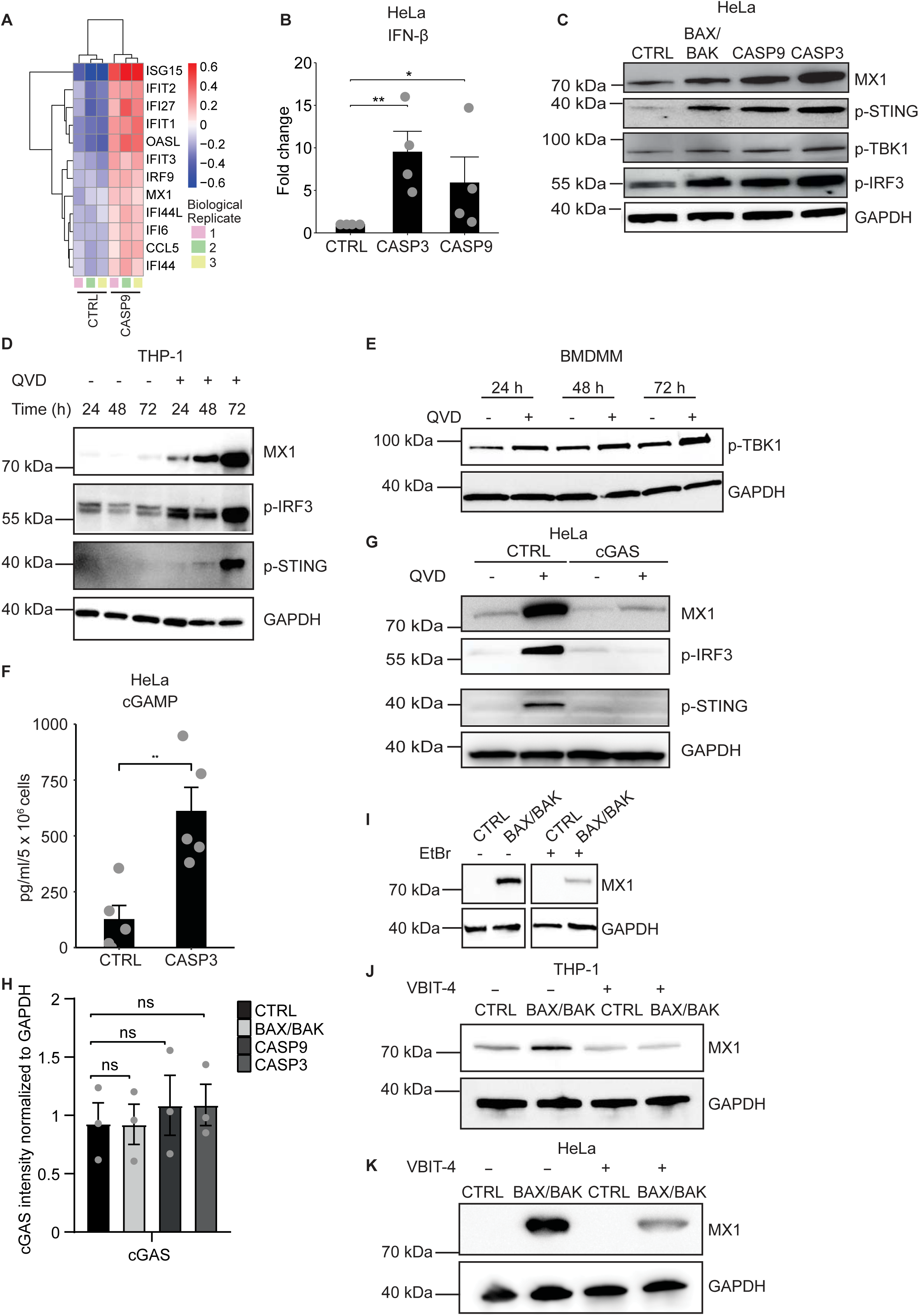
Constitutive activity of the mitochondrial apoptosis pathway downregulates cGAS activity and IFN signaling. **A** HeLa cell lines (CTRL, carrying a non-targeting gRNA or CASP9 CRISPR/Cas9 mutant) were cultured for 72 h followed by RNA isolation from lysates. RNA sequencing was performed. Heat map shows regulation of genes in the interferon signaling and response. **B** HeLa cell lines (CTRL, carrying a non-targeting gRNA or CRISPR/Cas9 mutants of the indicated genes) were evaluated for mRNA expression of IFN-β by RT-qPCR. Individual results are shown as dots and bars represent means of four individual experiments. Error bars show SEM. *, p < 0.05, **, p < 0.01. Significance was tested by one-way ANOVA with Dunnett’s post hoc testing. **C** Steady-state levels of MX1, p-STING, p-TBK1 and p-IRF3 in HeLa cells deficient in the mitochondrial apoptotic signaling components. GAPDH was used as a loading control. The result is representative of at least three individual experiments. **D** Undifferentiated THP-1 cells were treated with 10 µM QVD-OPh or DMSO (solvent control) for 24, 48 or 72 h. Levels of MX1, p-IRF3 and p-STING were analyzed by Western blotting. GAPDH was used as a loading control. The result is representative of three individual experiments. **E** BMDMM were differentiated with GM-CSF followed by 10 µM QVD-OPh treatment or DMSO (untreated, solvent control) for 24, 48 or 72 h. Post-treatment, levels of p-TBK1 were analyzed by Western blotting. GAPDH was used as a loading control. The result is representative of four individual experiments (cells from four individual mice). **F** Cells from HeLa cell lines (CTRL, carrying a non-targeting gRNA or CASP3 CRISPR/Cas9 mutant) were lysed after 24 h and steady-state cGAMP levels were measured by ELISA. Individual results are shown as dots and bars represent means of five individual experiments. Error bars show SEM. **, p < 0.01. The significance was tested by unpaired t-test. **G** Cells from HeLa cell lines (CTRL, carrying a non-targeting gRNA or cGAS CRISPR/Cas9 mutant) were treated with QVD-OPhor DMSO (solvent control) for 72 h. Levels of MX1, p-IRF3 and p-STING were analyzed by Western blotting. GAPDH was used as a loading control. The result is representative of at least three individual experiments. **H** Quantification of endogenous cGAS levels in HeLa cell lines (CTRL, carrying a non-targeting gRNA or indicated CRISPR/Cas9 mutants). For each condition, cGAS intensity was measured and normalized to the loading control (GAPDH) signal using ImageJ software. Individual results are shown as dots and bars represent means of three individual experiments. Error bars show SEM. Ns: p > 0.05. The significance was tested by one-way ANOVA with Dunnett’s multiple comparisons test. **I** Control or BAX/BAK-deficient HeLa cells were depleted of mtDNA HeLa cells by treatment with EtBr for 6 days, followed by recovery 48 h. Cells were lysed and levels of MX1 protein were analyzed by Western blotting. The figure shows results from the same membrane with lanes in between removed. **J** THP-1 cells were treated with 10 µM V-BIT4, harvested after 24 h and subjected to Western blotting. The result is representative of three individual experiments **K** HeLa cells were treated with 10 µM V-BIT4 for 72 h. Media supplemented with 10 µM V-BIT4 was changed each 24 h. After 72 h, cells were subjected to Western blotting. The result is representative of three individual experiments.

An increased IFN-response in cells lacking caspase-9 has been put down to the stimulation by mtDNA (*9*). Apoptotic release of mtDNA however required mitochondrial pores formed by BAX or BAK (*20–22*). We had found that the combined deletion of BAX and BAK in HeLa and THP-1 cells enhanced spontaneous cGAS signalling (see Fig. 2C, Fig. S3B, C). A stimulatory ligand can in this situation therefore not be released through BAX/BAK. We first tested whether mtDNA was required for the observed IFN response. Depleting BAX/BAK-deficient HeLa cells of mtDNA by prolonged culture in ethidium bromide reduced the MX1 levels (Fig. 2I), suggesting that mtDNA was required for its induction. It has also been reported that mtDNA can reach the cytosol through the voltage-dependent anion channel (VDAC), and the inhibitor of VDAC oligomerization, VBIT-4 (*23*), inhibited this release in published studies (*24, 25*). We found that treatment of THP-1 cells with VBIT-4 blocked the IFN-dependent MX1 induction caused by BAX/BAK deficiency (Fig.2J, Fig. S5) In HeLa cells, the IFN-inducing effect was also reduced although this was seen only after 72 h of treatment (Fig. 2K).

The results suggest that indeed mtDNA is the stimulus that drives spontaneous activation of cGAS and that is counter-acted by caspase-3. The release of mtDNA was BAX/BAK independent but appeared to depend on VDAC-oligomerization. Because cGAS is the receptor for mtDNA and cGAS activity was under the control of capase-3 (Fig. 2F), this indicates a direct effect of caspase-3 on cGAS. However, there was no detectable cleavage of cGAS in control HeLa cells, and no discernible difference in cGAS levels in cells lacking components of the mitochondrial apoptosis pathway (Fig. 2H). cGAS is an identified caspase-3-substrate (*10*), and it still seems plausible that it is direct caspase-3-mediated cleavage that may regulate cGAS activity but it cannot be by quantitatively relevant degradation as reported during apoptosis (*10*). Structural studies suggest that binding of cGAS to DNA causes conformational changes that may lead to dimerization of the protein (*26*) although we have been unable to detect cGAS dimers by cross-linking or on native gels. Indeed, it has already been reported that DNA-binding enhanced the cleavage of cGAS mediated by recombinant caspase-3 (*10*); caspase-3 therefore preferentially targets active cGAS over inactive cGAS. It seems likely that the low-level activity of caspase-3 found in non-apoptotic cells is only sufficient to cleave cGAS that has bound DNA – and is actively signalling –, and this portion is too small to be detected. In this way, caspase-3 would reduce spontaneous signalling of cGAS and limit the inflammation that would otherwise be caused by leakage of mtDNA into the cytosol. This mechanism would be similar to what we will show for MAVS in the following section.

### Blocking spontaneous, sub-lethal mitochondrial apoptosis signals enhances MAVS-signalling

The mitochondrial adapter of cytosolic RIG-I-like receptors (RIG-I and MDA5), MAVS, has also been identified as a caspase-3-substrate (*10*). We used transfection of HeLa cells with a ligand of RIG-I (5’-ppp-dsRNA) or of MDA5 (poly-I:C) to test for a regulation of MAVS signals by the apoptotic pathway. When HeLa cells were transfected with the RIG-I-agonist, the secretion of IL-6 was strongly enhanced in cells with deficiency of either BAX/BAK or caspase-3 (Fig. 3A, B; IL-6-secretion was completely MAVS-dependent, Fig. S6A). Phosphorylation of IRF3 by RIG-I ligand transfection, as well as the induction of MX1, were also enhanced in cells lacking these apoptosis signalling proteins (Fig. 3C). Enhanced IL-6 secretion was also seen upon transfection of poly-I:C into BAX/BAK-, caspase-3- or caspase-9-deficient cells (Fig. 3D-F).

**Fig. 3.**
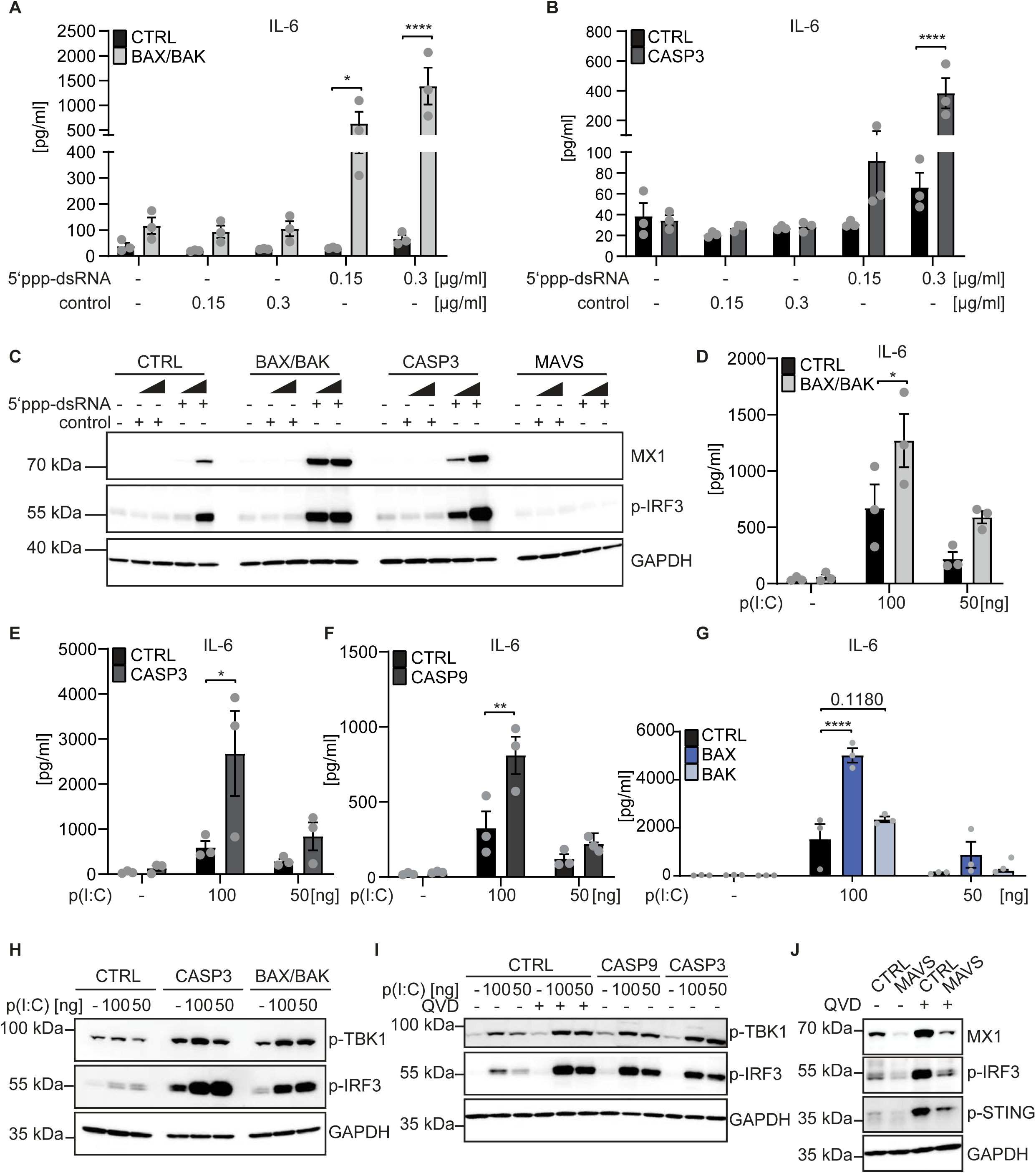
MAVS signals are enhanced by inhibition of the mitochondrial apoptosis pathway. Cells from HeLa cell lines (control or deficient in the indicated genes) were transfected with 0.15 µg/ml or 0.3 µg/ml 5’ppp-dsRNA (RIG-I-ligand) or control dsRNA (without 5’-triphosphate, non-stimulatory) **(A-C),** or transfected with 100 ng and 50 ng of poly (I:C). **A**, **B** At 24 h post-transfection, supernatants were harvested and subjected to IL-6 ELISA. Results from individual experiments are shown as symbols, and bars represent means of three individual experiments. Error bars show SEM and p values were calculated by two-way ANOVA (mean comparison) statistical assay with Sidak’s multiple comparisons testing. *, p < 0.05, ****, p < 0.0001. **C** Cells were transfected with RIG-I-ligand or non-stimulatory control dsRNA. 24 h post-transfection, cells were lysed and levels of MX1 and p-IRF3 were analyzed by Western blotting. GAPDH was used as a loading control. The result is representative of three individual experiments. **D-G** Cells were transfected with the indicated amount of poly (I:C). 24 h post-transfection, supernatants were harvested and subjected to IL-6 ELISA. Individual results are shown as symbols and bars represent means of three individual experiments. Error bars show SEM and p values were calculated by two-way ANOVA (mean comparison) statistical assay with Sidak’s multiple comparisons testing (D-F) and by two-way ANOVA (mean comparison) statistical assay with Dunnett’s multiple comparisons testing (G). *, p < 0.05, **, p < 0.01, ****, p < 0.0001. **H, I** Cells were pre-treated for 30 minutes with 10 µM QVD-OPh where indicated and transfected with the indicated amount of poly (I:C). 24 h post-transfection, cells were lysed and levels of p-TBK1 and p-IRF3 were analyzed by Western blotting. GAPDH was used as a loading control. The result is representative of at least three individual experiments. **J** HeLa cells (control or MAVS-deficient) were treated with QVD-OPh or DMSO (solvent control) for 72 h. Levels of MX1, p-IRF3 and p-STING were analyzed by Western blotting. GAPDH was used as a loading control. The result is representative of three individual experiments.

Phosphorylation of TBK1 and IRF3 was enhanced in poly-I:C-transfected cells deficient in BAX/BAK or caspase-3 (Fig. 3H), deficient in caspase-9 or treated with caspase-inhibitor (Fig. 3I). We confirmed the MAVS-dependency of IL-6 secretion and IRF3 phosphorylation during poly-I:C stimulation (Fig. S6B, C). There was minimal induction of cell death during these transfections (Fig. S7), indicating that the function of BAX/BAK and caspases was independent of apoptosis. The two effectors of mitochondrial apoptosis, BAX and BAK, are equivalent in many situations in that both can permeabilize mitochondria and cause apoptosis (*27*). Intriguingly, the effect we observed was almost entirely BAX dependent while BAK deficiency had no effect on IL-6-secretion during MAVS signalling (Fig. 3G). It therefore appears that the spontaneous activation of caspases is exclusively driven by BAX in these cells. Although the IFN response depended on cGAS/STING (see above), MAVS also made a contribution as the deletion of MAVS reduced MX1, pIRF3 and p-STING levels in steady state and when caspases were inhibited (Fig. 3J). These results argue for spontaneous signalling through MAVS, which also is under the control of constitutive caspase activity.

Infection with influenza A virus (IAV) is detected by RIG-I (*28*). IL-6 secretion as well as phosphorylation of IRF3 and TBK1 and induction of MX1 following IAV infection were strongly enhanced in BAX/BAK-deficient cells, compared to control cells (Fig. 4). The data show that MAVS-dependent signals received by cytosolic helicases are enhanced if sub-lethal signalling in the mitochondrial apoptosis pathway is blocked by deletion of any of the essential signalling proteins or by inhibition of caspases.

**Fig. 4.**
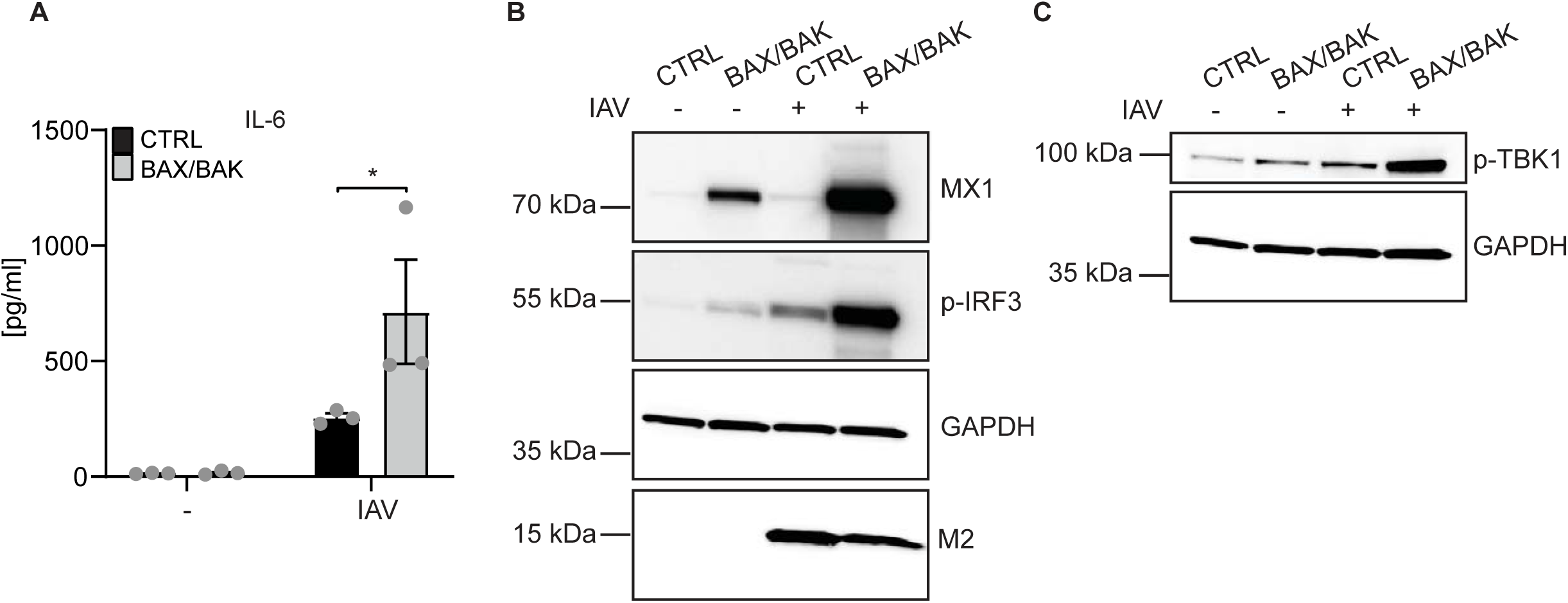
Loss of BAX/BAK enhances inflammatory signaling upon influenza A virus (IAV) infection. **A** Cells from HeLa cell lines (control or BAX/BAK-deficient) were infected with IAV (MOI of 1). 16 h post-infection, supernatants were harvested and subjected to IL-6 ELISA. Results from individual experiments are shown as symbols and bars represent means of three individual experiments. Error bars show SEM and p values were calculated by two-way ANOVA (mean comparison) statistical assay with Sidak’s multiple comparisons testing. *, p < 0.05. **B, C** Cells from HeLa cell lines (control or BAX/BAK-deficient) were infected with IAV (MOI of 1). 24 h post-infection, cells were lysed and levels of p-TBK1, MX1, p-IRF3 and IAV M2 protein were analyzed by Western blotting. GAPDH was used as a loading control. The result is representative of three individual experiments.

### Enhanced MAVS complex formation in the absence of spontaneous sub-lethal signalling

Upon ligand recognition, the cytosolic helicases translocate to intracellular membranes including mitochondria, where they induce the formation of MAVS oligomers (*29*). We used semi-denaturing detergent agarose gel electrophoresis (SDD-AGE) to detect MAVS signaling complexes. In unstimulated cells, a small amount of MAVS complexes was detected while the majority of the protein ran at a smaller molecular weight, presumably as monomers (Fig. 5A). Cells deficient in BAX/BAK, caspase-9 or caspase-3 all contained increased amounts of the higher molecular weight complexes, while the total amount of MAVS, as judged on SDS-PAGE, did not appear to be different (Fig. 5A, D). When MAVS was stimulated by transfection of poly-I:C, a discrete increase in signaling complexes was observed but this increase was strongly enhanced in cells deficient in BAX/BAK or deficient in caspase-3 (Fig. 5B). Treatment of control HeLa cells with caspase inhibitor also increased MAVS-containing complexes, similarly to the situation of BAX/BAK deficiency (Fig. 5C). In none of these situations, changes to the total amount of MAVS could be detected (Fig. 5A-D). Sub-lethal caspase activity that is generated by the mitochondrial apoptosis pathway therefore acts to decrease the amount of MAVS complexes that form spontaneously. This enhanced complex formation is a likely explanation for the enhanced MAVS signaling when these sub-lethal signals are blocked.

**Fig. 5.**
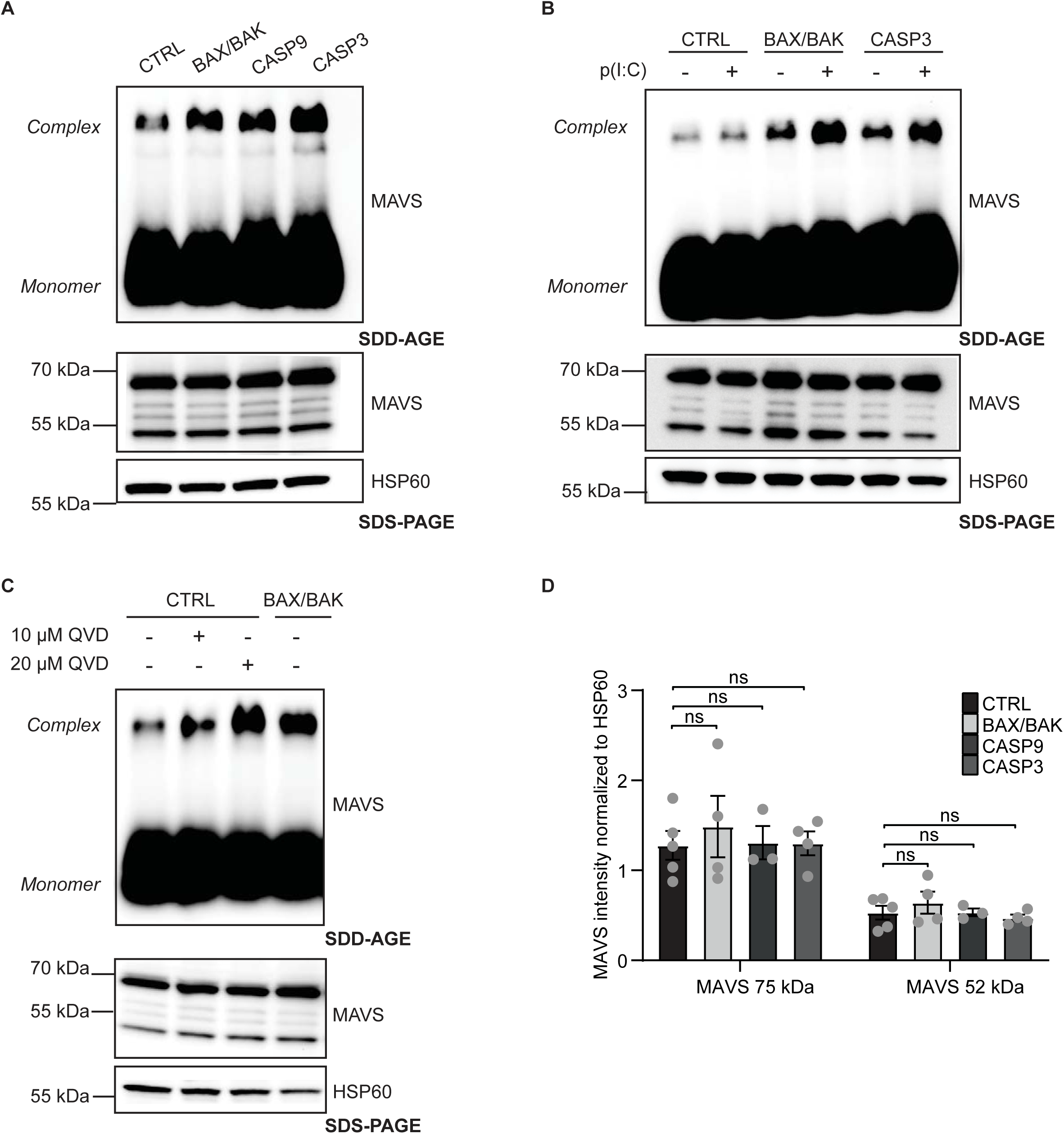
Levels of MAVS complexes but not of MAVS monomer are under the control of sub-lethal signals in the mitochondrial apoptosis pathway. **A** Heavy membrane fraction containing mitochondria were isolated from unstimulated HeLa cell lines (control or gene-deficient as indicated). Top panel, MAVS complexes and monomer were detected by SDD-AGE and Western blotting for MAVS. Bottom panel, aliquots of the same fractions were analyzed by SDS PAGE. The mitochondrial marker HSP60 was used as a loading control. The results are representative of at least three individual experiments. **B** Cells from HeLa cell lines (control or gene-deficient as indicated) were mock transfected or transfected with 1.3 µg poly (I:C). At 24 h post-transfection, heavy membrane fractions were isolated and analyzed by SDD-AGE or SDS-PAGE as in A. The results are representative of at least three individual experiments. **C** Cells from HeLa cell lines (control or gene-deficient as indicated) were treated with the indicated concentrations of QVD-OPh for 16 h. Heavy membrane fractions were isolated and analyzed by SDD-AGE or SDS-PAGE as in A. The result is representative of at least three individual experiments. **D** Quantification of the endogenous MAVS levels from SDS-PAGE from experiments as shown in A. Bands for the two MAVS isoforms were quantified from Western blots of SDS gels and were normalized to HSP60. Results from individual experiments are shown as symbols and bars represent means of five (control), four (BAX/BAK- and caspase-3-deficient cells) or three (caspase-9-deficient cells) individual experiments. Error bars show SEM and p values were calculated by two-way ANOVA (mean comparison) statistical assay with Sidak’s multiple comparisons testing. Ns: p > 0.05.

### The MAVS complex has higher sensitivity to caspase-3 than the monomer

These results suggested a scenario where low-level caspase-3 cleaved only the MAVS complex but not the monomer, suggesting higher sensitivity of the complexes to proteolysis by caspase-3. To test this directly, we isolated heavy membrane fractions from BAX/BAK-deficient cells (which contain a high amount of spontaneously forming MAVS complexes) and digested them with graded concentrations of recombinant caspase-3. At a concentration of 0.5 ng/µl, no degradation of MAVS monomers was detected. However, the complexes already showed significant proteolytic degradation at this concentration of caspase-3 (Fig. 6A, C). Higher concentrations of the protease degraded both complexes and monomers but the sensitivity remained clearly higher in the complex fraction. Because we observed some background degradation without addition of the recombinant enzyme (not shown), which may have been the result of co-purification of caspase-3 from the extracts, we repeated the experiments with heavy membrane fractions from caspase-3-deficient cells. We detected the same pattern of enhanced degradation of MAVS complexes compared to the monomeric form as in fractions from BAX/BAK-deficient cells (Fig. 6B, D).

**Fig. 6.**
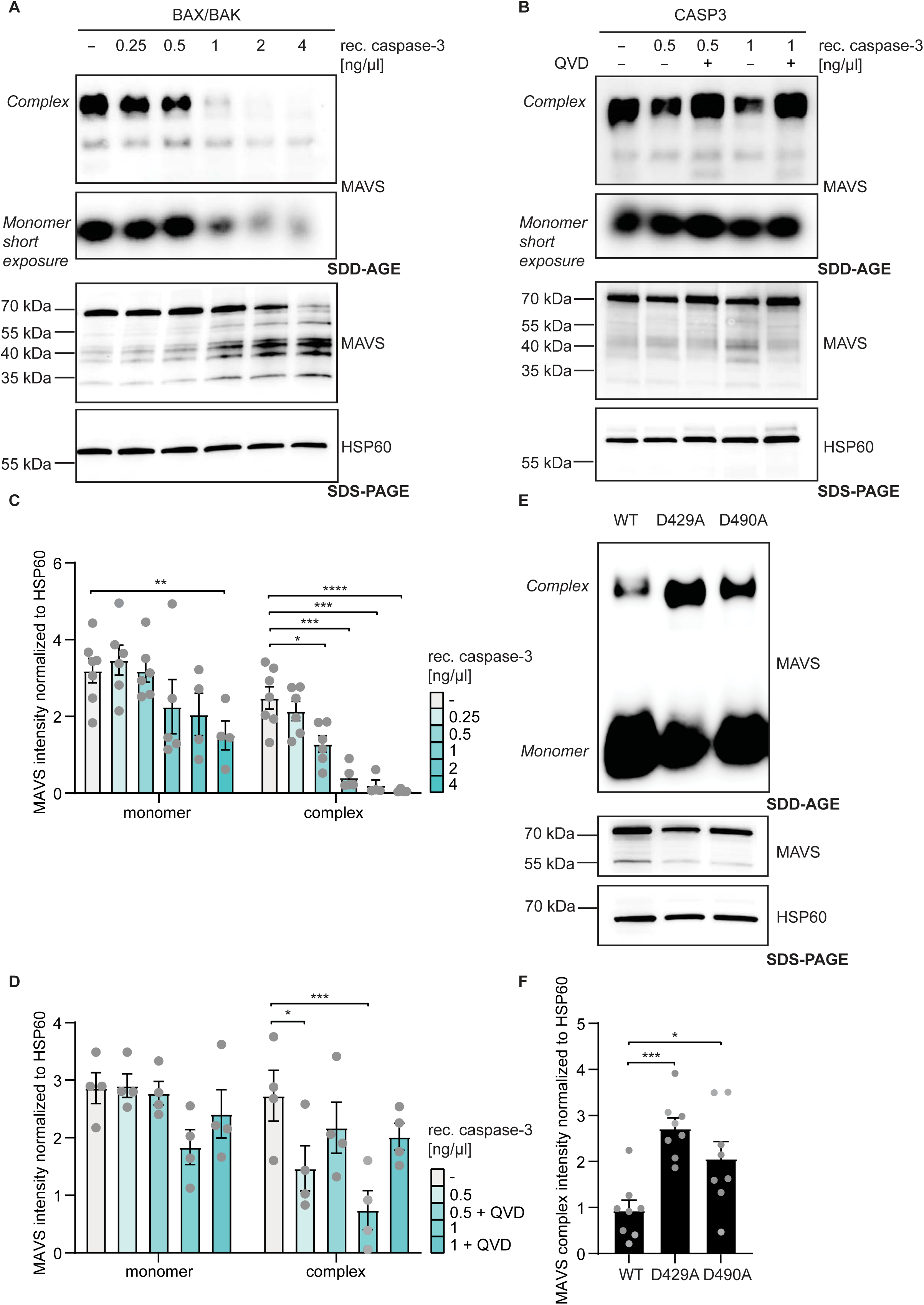
Differential sensitivity of MAVS complexes vs. monomer to caspase-3-mediated proteolysis. **A, B** Heavy membrane fractions were isolated from HeLa cell lines (BAX/BAK- or caspase-3-deficient). The fractions were incubated with the indicated concentrations of recombinant caspase-3. To some aliquots, 10 µM QVD-OPh was added prior to caspase-3 addition (B). Samples were incubated at 37°C for 30 min. Reactions were stopped by addition of 10 µM QVD-OPh. Reactions were loaded on SDD-AGE and SDS-PAGE. HSP60 was used as a loading control. The result is representative of at least three individual experiments (quantified in C, D). **C, D** Quantification of MAVS monomer and complex signals from SDD-AGE, normalized to HSP60 signal (SDS-PAGE) on Western blots as in A and B (C refers to A, D to B). Results from individual experiments are shown as symbols, and bars represent means of at least four individual experiments. Error bars show SEM and p values were calculated by two-way ANOVA (mean comparison) statistical assay with Dunnett’s multiple comparisons testing. Ns: p > 0.05, *, p < 0.05, **, p < 0.01, ***, p < 0.001, ****, p < 0.0001. **E** Heavy membrane fractions were isolated from unstimulated HeLa cell lines (controls or mutants (single clones) with a mutation of either of the two caspase-3 cleavage sites D429A or D490A. Levels of MAVS complexes and monomer were analyzed as in A. HSP60 was used as a loading control for crude mitochondrial fraction. The result is representative of at least eight individual experiments. **F** Quantification of MAVS complex signals from SDD-AGE, normalized to HSP60 signal (SDS-PAGE) on Western blots as shown in E. Results from individual experiments are given as symbols, and bars represent means of eight individual experiments. Error bars show SEM and p values were calculated by one-way ANOVA (mean comparison) statistical assay with Dunnett’s multiple comparisons testing. *, p, ***, p < 0.001.

MAVS has two caspase-3 cleavage site that have been mapped (*10*). We mutated endogenous MAVS in the genome of HeLa cells to generate the two separate mutations D429A and D490A, destroying either of the two caspase-3 cleavage sites. We selected individual clones carrying these mutations. As shown in Fig. 6E, F, these clones had substantially higher levels of MAVS complexes in steady state, confirming the hypothesis that caspase-3 cleavage of MAVS is the mechanism of reducing the complexes. These results support a model where MAVS complexes form in human cells in the absence of specific stimulation, probably in response to the cytosolic presence of dsRNA. Low-level, sub-lethal caspase-3 activity that is generated through activity of the mitochondrial apoptosis pathway, specifically cleaves the signaling complexes and reduces signals generated in steady state.

We finally tested whether IRF3, the third identified substrate of caspase-3 in innate immunity, also follows the pattern of preferential degradation of the protein complex. However, this did not seem to be the case. IRF3 is phosphorylated and forms dimers upon its activation. Treatment of HeLa cells with caspase inhibitor increased the level of IRF3 dimer (Fig. S8A, B). However, the sensitivity of dimer and monomer to proteolysis by recombinant caspase-3 was similar (Fig. S8C, D). This suggests that the increase in IRF3 dimer formation upon caspase inhibition is the result of increased upstream signaling input, most likely from cGAS and perhaps MAVS, rather than a direct effect of caspase-3 on IRF3.

## Discussion

Caspases block inflammation by cleaving signalling proteins of innate immunity. During apoptosis, this appears to involve the quantitative degradation of cGAS, MAVS and IRF3, to make them unavailable for signalling. We here illustrate a novel concept: the activity of caspase-3 is higher towards activated than inactive forms of these proteins. Caspase-3 appears to be constitutively active through basal activity of the mitochondrial apoptosis pathway in all cells tested. This activity acts to tune the responsiveness of the cells to infection and may have the function of blocking signals that would be constitutively generated by leakage of mitochondrial constituents.

In the case of cGAS, the enhanced sensitivity of the protein is probably generated by binding of stimulatory DNA, as has indeed been reported before with recombinant caspase-3 (*10*). Ligand binding causes a conformational change in cGAS and is predicted to induce cGAS complex formation (*26*). We have been unable to detect cGAS complexes in native gels or by chemical cross-linking. Mutation of the caspase-3 cleavage site in cGAS inactivates the enzyme (*10*). We reproduced this finding: genomic mutation of D319 blocked IFN signalling in the cells (not shown); we therefore were unable to test the effect of the mutation functionally. Caspase-3 very likely acts directly on cGAS because deletion of caspase-3 enhanced cGAS activity, and cGAS is the receptor for mtDNA. While thus final proof is lacking, the data strongly suggest that specific caspase-3-mediated cleavage of DNA-bound, activated cGAS reduces spontaneous IFN I signalling in human cells. Inactive cGAS however remains unaffected, explaining how cGAS cleavage by caspase-3 is required to reduce the basic IFN signal but the total level of cGAS is not detectably reduced.

We were however able to test the concept for MAVS. MAVS signalling complexes were increased in the absence of activity of the mitochondrial apoptosis pathway and were more sensitive to caspase-3 cleavage. Mutation of either of the two caspase-3 cleavage sites led to accumulation of MAVS complexes. The data suggest that spontaneous signals through MAVS drive its activation while caspase-3 specifically degrades MAVS complexes. Although these signals were insufficient to lead to IL-6 secretion, they appear to be sufficient to induce some level of MAVS complex accumulation and contribute to spontaneous IFN production. At least as a monomer, MAVS is probably largely intrinsically unstructured (AlphaFold prediction, https://www.uniprot.org/uniprotkb/7Z434/entry#structure), and the caspase-3 cleavage sites are located in the unstructured regions. Activation of MAVS involves binding of both protein and RNA (*30*) and very likely induces secondary structures in the protein, which may stabilize the caspase cleavage sites and make them more easily accessible to the protease.

It was surprising to note that in addition to caspase-9, only caspase-3 but not caspase-7 is active in HeLa cells. We have reported that the mitochondrial apoptosis pathway is constitutively active at a sub-lethal level in all tested cells, including mouse embryonic fibroblasts and mouse intestinal organoids, and it drives growth behaviour through the activation of the caspase-activated DNase (*18*). During apoptosis, caspase-9 activates both caspase-3 and caspase-7. Both effector caspases have almost identical cleavage preferences towards peptide substrate (*11*) although some differences in function appear to exist (*12*). Both caspase-3 and -7 are activated by caspase-9 dependent cleavage between the small and large subunits, and this cleavage site in caspase-3 fits the (peptide) substrate preference of caspase-9 better than the site in caspase-7 (*11*). It is therefore not unlikely that a higher level of active caspase-9 is required for the activation of caspase-7 than of caspase-3, and this may not be achieved in a sub-lethal setting.

In the cells tested, there was constant activation of cGAS that was counter-regulated by caspase-3. The data suggest that mtDNA leaks out of mitochondria even without experimental manipulation of the cells. This availability of cytosolic mtDNA did not depend on BAX/BAK pores but the inhibitor of VDAC oligomerization VBIT-4 blocked the signal, suggesting that spontaneous, VDAC-mediated release of mtDNA into the cytosol occurs. The results suggest that activation of caspase-3 has a function in reducing such spontaneous cGAS signals and inflammation in the absence of infection. Spontaneous MAVS-dependent downstream signalling was marginal in our cells, although spontaneous complex formation and a contribution to IFN signalling indicates the presence of a signal. Cytosolic dsRNA, presumably of mitochondrial origin, has been identified as one driver of the senescence associated secretory phenotype (*31*) [preprint]. Although senescence-associated stress enhanced the translocation of dsRNA, it was also detected in normally proliferating, non-senescent fibroblasts in that study. Caspase-3 activation, and caspase-3-dependent cleavage of MAVS, may therefore be required to reduce dsRNA-induced inflammation at least in situations of cellular stress but probably also in resting cells.

Our results identify an interesting principle, that caspase-sensitivity of substrate proteins changes dependent on the activation status of the protein. While the data demonstrate this for MAVS and strongly suggest it for cGAS, there is the potential of a broader application of this principle. Of the many hundreds of caspase-3 substrates identified, a convincing physiological role as a substrate has been described for very few. It may be worth considering caspase-3-dependent cleavage of a small part of the protein as a regulatory mechanism, not exclusively of total substrate levels but of the conformational, functional state of target proteins.

## Methods

### Cell culture

HeLa cells (ATCC, cat. CCL-2.1) were cultured as described previously (*32*). THP-1 cells were cultured in RPMI 1640 (PAN-Biotech, cat. P04-17525) w/o: L-Glutamine, w: 2.0 g/L NaHCO3, Very Low Endotoxin, 10% hi (heat-inactivated) FCS, 1% p/s (penicillin/streptomycin) and 2 mM GlutaMAX™ Supplement (Thermo Fisher Scientific, cat. 35050061). Cells were incubated at 37°C in 5% CO_2_. Primary Bone Marrow-Derived Monocytic Macrophages (BMDMM) were prepared as described (*33*). 4 × 10^5^ cells/well were seeded in a 24-well non-tissue culture-treated plate in a media containing recombinant GM-CSF (20 ng/ml). 4 h post-seeding, cells were treated for 24, 48 and 72 h with 10 µM QVD-OPh were lysed in 100 µl 1× SDS-Laemmli buffer, boiled at 95°C and subjected to Western blotting. Gene-deficient cell lines were generated using the lentiCRISPRv2_puro system (*34*) as described (*32*). HeLa cells: control, STING-, Caspase-9, -3, -7, BAX/BAK and BAX or BAK-deficient cells have been described (*32, 35–37*). The HeLa MAVS-deficient cell line used here was selected from a polyclonal line (*32*). cGAS-deficient HeLa cells were generated with the gRNA ATGATATCTCCACGGCGGCG. We further generated a different cGAS deletion mutation using recombinant Cas9 protein with a chemically modified gRNA (5’-AATATCTGTGGATATAACCC-3’; designed using the Alt-R™ CRISPR HDR Design Tool (IDT, Integrated DNA Technologies)). The same gRNAs were used for gene-deficient and control THP-1 cells (*35*). For MAVS point mutations at the endogenous locus, we used the Alt-R™ CRISPR HDR Design Tool (IDT, Integrated DNA Technologies). Sequences for HDR donor oligonucleotides (-strand) to generate D429A and D490A mutations at the MAVS locus are available upon request.

### Enzymatic caspase-3 like cleavage assay (DEVD-AMC cleavage assay)

Cells were lysed in 1x lysis buffer (Cell Signaling, cat. 9803) supplemented with protease inhibitor (cOmplete EDTA-free Protease Inhibitor Coctail Roche, Sigma-Aldrich, cat. 4693132001). 10 µl of the lysate was incubated with 90 µl of reaction buffer (MDB buffer (10 mM HEPES, pH 7, 40 mM β-glycerophosphate, 50 mM NaCl, 2 mM MgCl_2_, and 5 mM EGTA) supplemented with 100 µg/ml BSA, 0.1% CHAPS, and 11 µM Ac-DEVD-AMC) in triplicates. Fluorescence was measured using a Spark 10M microplate reader (Tecan) (380/460 nm. The fluorescence readings were normalized to the protein concentration (Bradford).

### N-terminomics with 16-plex tandem mass tag (TMT)

HeLa cells were cultured for 24 h. Cells were lysed, reduced and alkylated at 95 °C for 10 min in lysis buffer (6 M GuHCl, 10 mM TCEP, 40 mM CAA, 200 mM HEPPS, pH 8.5), followed by sonication and protein quantification. SP3 cleanup was performed using 10 μL of SP3 beads (1:1 Sera-Mag Speed Beads A and B) in 80% EtOH, incubated for 18 min at 800 rpm, magnetically immobilized and washed twice with 80% EtOH and once with ACN. Proteins were resolubilized in 2 M GuHCl, 10 mM TCEP, 200 mM HEPPS (pH 8.5) and labelled with TMT pro 16-plex reagents (*38*) for 1 hour at 20 °C, followed by quenching with hydroxylamine. Labelled samples were combined, SP3 cleanup was performed followed by digestion with trypsin (1:50, w/w) overnight at 37°C in 100 mM HEPES (pH 7.6). To remove internal peptides, the digested samples were incubated overnight with 30 mM sodium cyanoborohydride and an amine-reactive aldehyde-derivatized polymer (*39*) (1:5, w/w peptide:polymer), quenched with 100 mM Tris, filtered (Amicon 30 kDa), acidified (1% FA), desalted, and fractionated into six high pH RP fractions (*40*). Peptides were analyzed on a Dionex Ultimate 3000 UHPLC+ coupled to an Orbitrap Eclipse mass spectrometer, separated on a C18 analytical column (75 μm × 45 cm, 3 μm) using an 80-min gradient (8–34% ACN, 0.1% FA, 5% DMSO). The Orbitrap operated in data-dependent acquisition (DDA) with MS1 scans (m/z 360–1500, 60K resolution), MS2 scans (15K, HCD NCE 34%), and MS3-based TMT quantification (30K, NCE 55%). Dynamic exclusion was 90s. Raw MS data were analyzed using FragPipe (v20.0) with MSFragger (v3.8) (*41*), searching against the human UniProtKB database (07.2021) with a TMT16-MS3 workflow (20 ppm precursor tolerance, semi-tryptic specificity). Perseus (v2.0.10.0) was used for data analysis (*42*). Differential analysis (CTRL vs caspase deficient cell line) was conducted using a Student’s t-test (S0 = 0.1, FDR = 5%). MS data are available via ProteomeXchange PRIDE (*43*), dataset PXD058771.

### RNA isolation and RT-qPCR

Samples were lysed in TRI Reagent and RNA was extracted using the Direct-zol RNA MicroPrep or RNA Miniprep kit (Zymo Research, cat. R2061/R2053). RNA purity and concentration were assessed using a NanoDrop spectrophotometer. RNA was reverse transcribed using RevertAid First Strand cDNA synthesis kit (Thermo Fisher Scientific, cat. K1622). For qPCR, reactions were prepared in a 384-well plate using 1X SYBR Green master mix (Applied Biosystems, cat. 4472918). Results were analyzed using QuantStudio Design and Analysis software v.1.5.2. Primer sequences are available upon request. The comparative CT method (ΔΔCT method) was used to determine relative expression of target gene to the housekeeping gene.

### Bulk RNA sequencing and analysis

Library preparation for bulk-sequencing of poly(A)-RNA was done as described previously (*44*). Barcoded cDNA of each sample was generated with a Maxima RT polymerase (Thermo Fisher) using oligo-dT primer containing barcodes, unique molecular identifiers (UMIs) and an adaptor. 5’-Ends of the cDNAs were extended by a template switch oligo (TSO) and full-length cDNA was amplified with primers binding to the TSO-site and the adaptor. NEB UltraII FS kit was used to fragment cDNA. After end repair and A-tailing a TruSeq adapter was ligated and 3’-end-fragments were finally amplified using primers with Illumina P5 and P7 overhangs. The library was sequenced on a NextSeq 500 (Illumina) with 61 cycles for the cDNA in read1 and 19 cycles for the barcodes and UMIs in read2. Data were processed using the published Drop-seq pipeline (v1.0) to generate sample- and gene-wise UMI tables (*45*). Reference genome (GRCh38) was used for alignment. Transcript and gene definitions were used according to GENCODE v38. Data analysis was done with R. Package ‘apeglm’ for LFC shrinkage (*46*). The analysis of differentially expressed genes was performed with DESeq2 package (*47*). A log fold change threshold was set at ±0.58 (approximately 1.5-fold change) and a p-value threshold of 0.05, adjusted for multiple testing using the Benjamini-Hochberg method. For enrichment analysis, reactome pathway analysis was performed using the ReactomePA package (*48*).

### ELISA

The levels of 2′,3′-cGAMP in total cell lysates were quantified using the DetectX 2′,3′-Cyclic GAMP Enzyme Immunoassay Kit (Arbor Assays, K067-H1). In brief, 5 × 10^6^ cells were trypsinized, counted, and lysed in lysis buffer. The lysates were further processed by sonication as previously described (*49*). IL-6 levels were detected from supernatants after indicated treatments with human IL-6 ELISA MAX Deluxe kit (BioLegend, cat. 430516).

### Western blotting

Cells were lysed in the plates in 1× SDS-Laemmli buffer or RIPA buffer, boiled and run on SDS-PAGE. Proteins were transfer to PVDF or nitrocellulose membranes. Membranes were incubated with the primary antibodies: MX1 (Clone M143, provided by G. Kochs, Virology Freiburg), p-STING (Cell Signaling, cat. 19781), p-TBK1 (Cell Signaling, cat. 5483), p-IRF3 (abcam, cat. ab76493), GAPDH (Millipore, cat. MAB374), MAVS (Proteintech, cat. 14341-1-AP), HSP60 (Cell Signaling, cat. 4870), cGAS (Cell Signaling, cat. 15102), IRF3 (IBL-America, cat. 18781), BAX (Cell Signaling, cat. 2774S), BAK (Calbiochem, cat. AM03) or IAV M2 protein (Thermo Fisher Scientific, cat. MA1-082).

### mtDNA depletion

1.0 × 10^5^ HeLa cells were seeded in the 6-well plate. The next day, media was replaced to regular cell culture media supplemented with sodium pyruvate (100 µg/ml) (Gibco, cat. 11360-039) and uridine (50 µg/ml) (Sigma-Aldrich, cat. U3003-5G). To deplete mtDNA cells were treated with ethidium bromide (EtBr) (50 ng/ml) (Thermo Fisher Scientific, cat. 17896) as reported (*50*). Six days later, EtBr was removed and cells were allowed to recover for 48 h before analysis.

### Transfection with 5‘ppp-dsRNA (RIG-I-ligand) or poly (I:C), other reagents

1.5 × 10^5^ HeLa cells were seeded in the 6-well plate. The next day, cells were transfected with 5’ppp-dsRNA (RIG-I-ligand; Invivogen, cat. tlrl-3prna), 5’ppp-dsRNA control (Invivogen, cat. tlrl-3prnac; dsRNA without 5’-triphosphate) or 100 ng poly (I:C) (final concentration in the well 60 ng/ml) or 50 ng poly (I:C) ((Invivogen, catalog no. tlrl-pic-5); final concentration in the well 30 ng/ml). dsRNA or sterile endotoxin-free water (referred as untreated) and lipofectamine (RNAiMAX transfection reagent; Invitrogen, Thermo Fisher Scientific; cat. 13778030) were first diluted in OPTIMEM medium (Invitrogen, Thermo Fisher Scientific, cat. 11058021) and mixed together in ratio 1:2 (µg of dsRNA : µl of lipofectamine). Mixtures were incubated for 30 min at RT and finally added to the cells. Cell death measurement was done with combined supernatant and trypsinized cells. diABZI compound 3 (STING agonist; Invivogen, cat. tlrl-diabzi), QVD-OPh from Hycultec (cat. HY12305; 25 mg), V-BIT4 (Selleck Chemicals, cat. S3544) and recombinant human caspase-3 (R&D Systems, cat. 707-C3-010/CF) were used as indicated in the figure legends. VBIT-4 was used at 10 µM for 24 (THP-1) or 72 (HeLa cells) hours.

### IAV infection

Influenza A virus (IAV) (A/WSN/1933 - H1N1) was kindly provided by Georg Kochs, Institute of Virology, Freiburg. 3 × 10^5^ HeLa cells were seeded in a 6-well plate the day before infection. Cells were infected at MOI=1 as described (*51*).

### Mitochondrial isolation and Semi-denaturing detergent agarose gel electrophoresis (SDD-AGE)

5 × 10^6^ cells were seeded in 15 cm plates the day before treatment. Cells were transfected with poly (I:C) (final concentration in the plate 60 ng/ml) and harvested 24 h post-treatment. 4.5 × 10^6^ cells were seeded in 15 cm plates the day before treatment. Some aliquots were pretreated with QVD-OPh for 16 h. Cells were suspended in MB-EDTA buffer (210 mM mannitol, 70 mM sucrose, 10 mM HEPES pH 7.5, 1 mM EDTA pH 7.5, supplemented with 1x protease inhibitor (cOmplete EDTA-free Protease Inhibitor Coctail Roche, Sigma-Aldrich, cat. 4693132001) and 1x phosphatase inhibitors (PhosSTOP, Roche, Sigma-Aldrich, cat.4906845001)). Heavy membrane fractions containing mitochondria were obtained by passing cells through a 27 G needle, followed by 10 000 x g centrifugation at 4°C, as previously described (*52*). 25 µg protein (Bradford) were mixed with SDD-AGE sample buffer to achieve 1x concentration (0.5 x TAE, 5% glycerol, 2% SDS, bromphenol blue) and incubated for 5-10 min at RT. SDD-AGE samples were run on the 1.5% agarose gel with 0.1% SDS, in running buffer (1x TAE with 0.1% SDS) at 30 V for 3-4 hours, followed by capillary transfer (*53*). 25 µg were also used for SDS-PAGE. MAVS was detected as above.

### MAVS cleavage by recombinant caspase-3

Heavy membrane fractions were isolated and pelleted at 10 000 x g at 4°C. Pellets were resuspended in a freshly prepared reaction buffer (25 mM HEPES pH 7.5, 0.1% CHAPS, 10 mM DTT). 25 µg protein were mixed with recombinant caspase-3 (diluted in the reaction buffer). Samples were mixed gently and incubated at 37°C for 30 min. Reactions were stopped by addition of 10 µM QVD-OPh. 15 µg of protein were loaded on SDD-AGE or SDS-PAGE.

### Quantification of signals

Bands were quantified using ImageJ. Statistical analysis was done as indicated in the figure legend.

### Annexin V/PI staining

Cells were incubated with Annexin V-FITC (Invitrogen, cat. BMS306FI-300) in binding buffer (BD Pharmingen, cat. 556454). Propidium iodide (PI) (Sigma-Aldrich, catalog no. P4170) was added to the cell suspension. Samples were analyzed on a FACSCalibur (Becton, Dickinson).

### IRF3 dimerization and caspase-3-cleavage

Dimerization of IRF3 was analyzed by native-PAGE as described (*54*). Aliquots of the same fractions were also run on SDS-PAGE. Whole cell extracts were incubated with recombinant caspase-3 (diluted in reaction buffer) at 37 °C for 1 h. Samples were run on native PAGE and SDS-PAGE. Bands were quantified using ImageJ and normalized to GAPDH intensity from SDS-PAGE blots.

## Acknowledgement and Grant information

This work was supported by the Deutsche Forschungsgemeinschaft (DFG), GRK 2606 (ProthPath) to G.H. We acknowledge funding from the EU-H2020-MSCA-COFUND EURIdoc programme (No. 101034170) (to G.H.). The Orbitrap Eclipse mass spectrometer was funded in part by the German Research Foundation (INST 95/1650-1). We thank Juliane Vier for help in modifying THP-1 cells, Abdul Moeed for IAV stock preparation and Georg Kochs for antibodies.

## Legends to supplemental figures

**Fig. S1.**
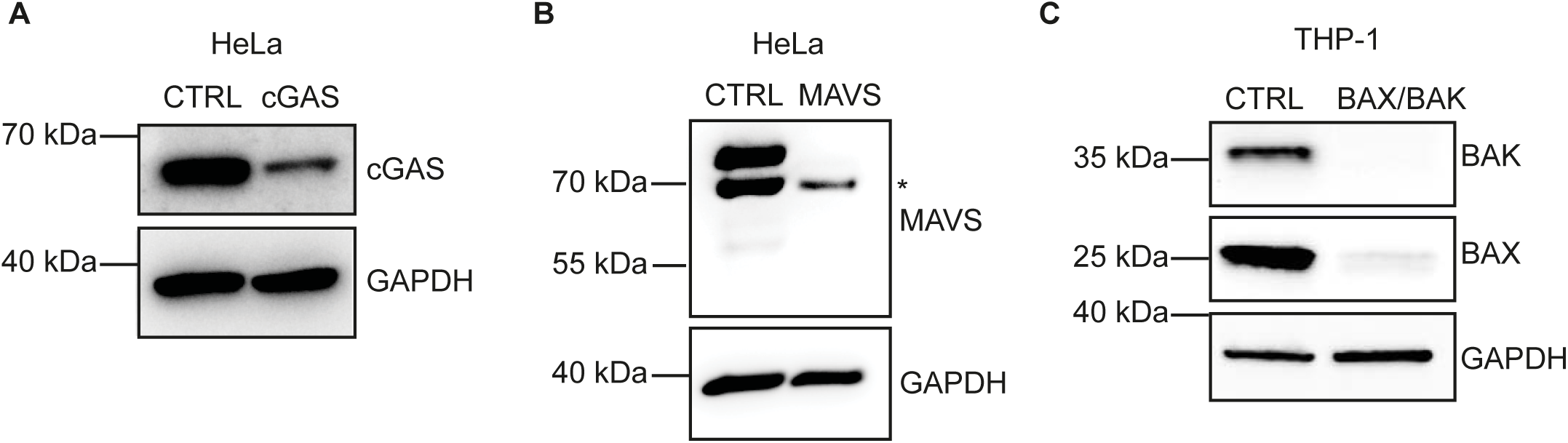
Western blots confirming protein deficiency in the lentiCRISPRv2 cell lines used in the study. **B,** Asterisk indicates a non-specific band.

**Fig. S2.**
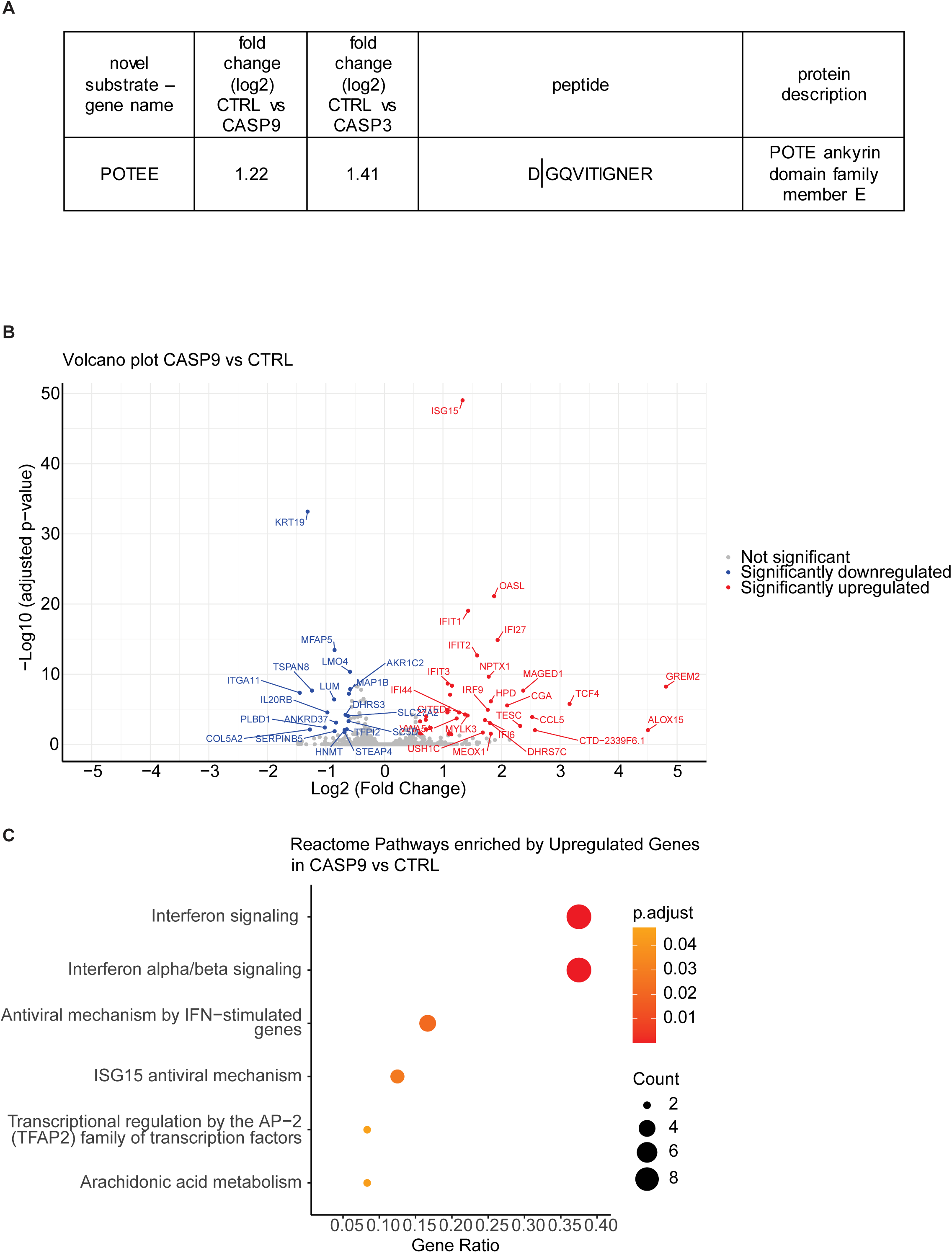
Caspase-9 and -3 activity acts to downregulate a spontaneous IFN response. **A** A novel putative caspase substrate was identified by N-terminomics. N-termini appears significantly less abundant in the caspase-9-deficient cells. The identified peptide is shown. **B** Volcano plot showing genes with differential expression in HeLa CTRL cells compared to CASP9 CRISPR/Cas9 mutant. Experiment was done three times. The criteria for differentially expressed genes were a log2-fold change ≥ 0.58 or ≤ −0.58 and an adjusted p value ≤ 0.05. **C** Reactome pathway database analysis showing pathways defined by genes upregulated in steady state of caspase-9-deficient vs. control HeLa cells. The data are the result from three independent experiments. Significance, gene count and gene ratio classified within each pathway are shown. Count represents the number of up-regulated genes associated with each pathway. Gene ratio represents proportion of up-regulated genes associated with each pathway relative to the total number of up-regulated genes in the data set.

**Fig. S3.**
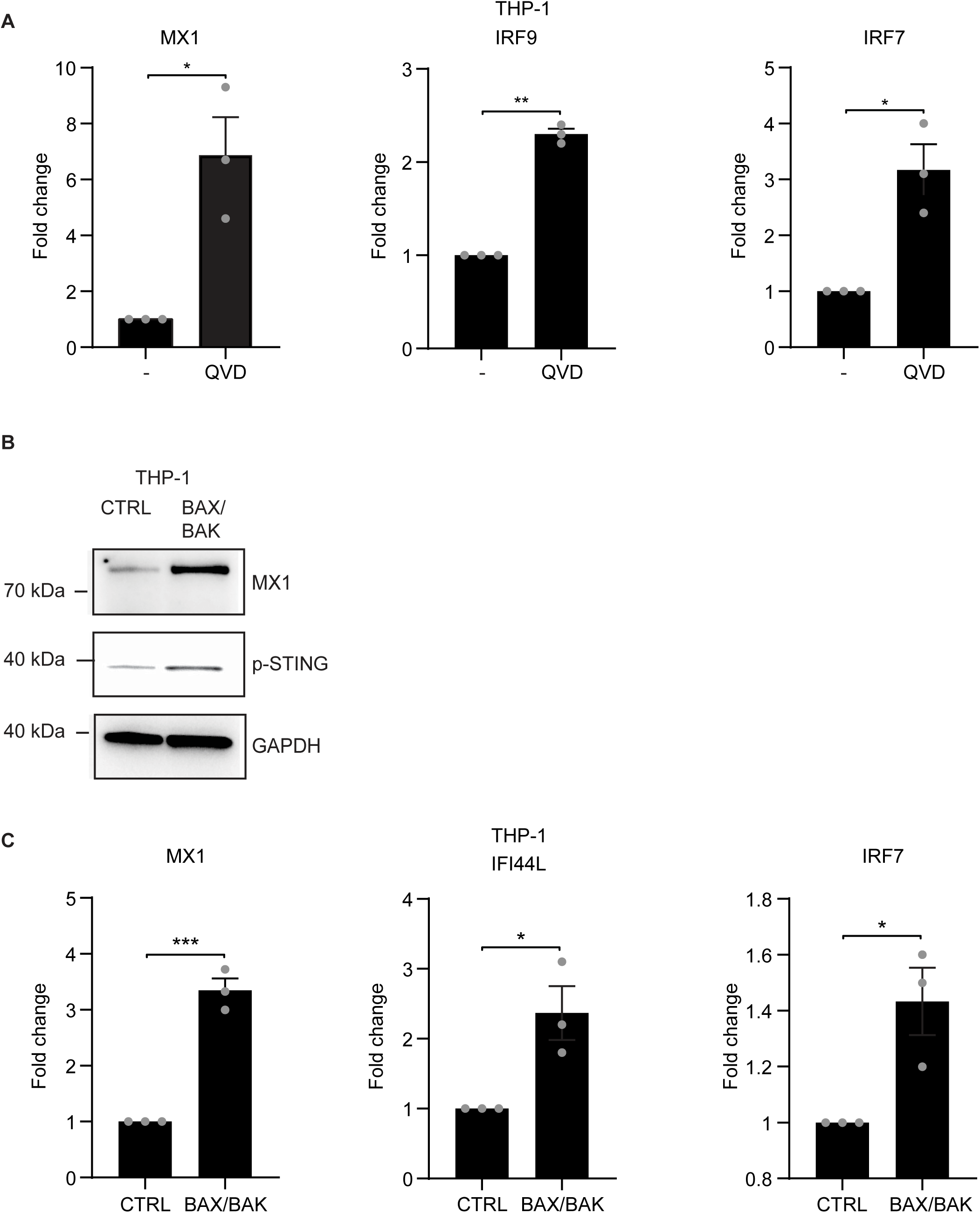
IFN response induced by inhibition of caspase-signaling. **A** THP-1 cells were treated with 10 µM QVD-OPh or DMSO (solvent control) for 8 h and were evaluated for mRNA expression of MX1, IRF9 and IRF7 by RT-qPCR. Individual results are shown as dots and bars represent means of three individual experiments. Error bars show SEM. The significance was tested by paired t-test. *, p < 0.05, **, p < 0.01. **B** Steady-state levels of MX1 and p-STING were evaluated in THP-1 cells (control or BAX/BAK-deficient). GAPDH was used as a loading control. The result is representative of three individual experiments. **C** THP-1 cells (control of BAX/BAK-deficient) were evaluated for mRNA expression of MX1, IFI44L and IRF7 by RT-qPCR. Individual results are shown as dots and bars represent means of three individual experiments. Error bars show SEM. The significance was tested by unpaired t-test. *, p < 0.05, ***, p < 0.001.

**Fig. S4.**
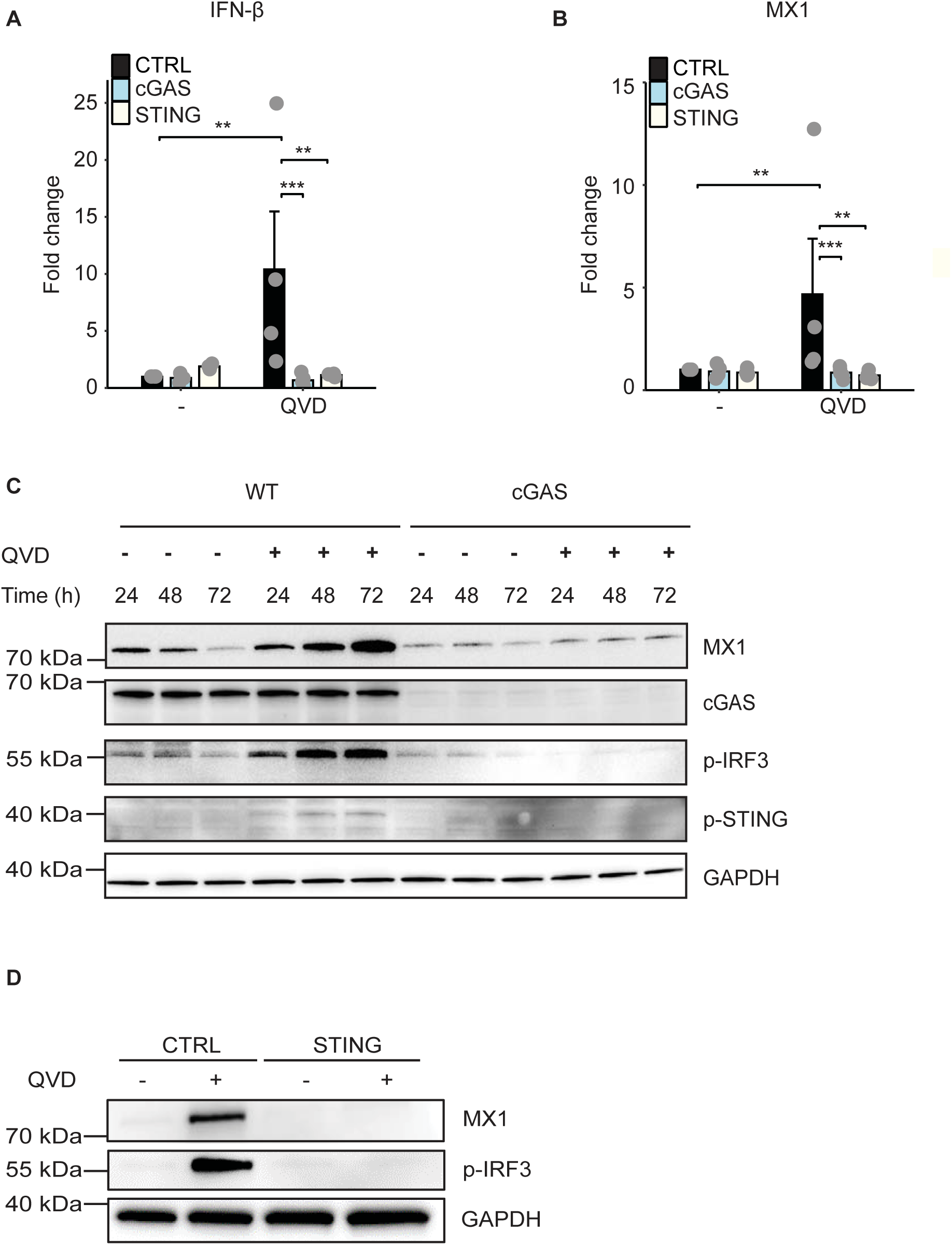
cGAS and STING dependency of caspase control of IFN signaling. **A, B** Cells from HeLa cell lines (control or cGAS- or STING-deficient) were cultured for 56 h with 10 µM QVD-OPh or DMSO (solvent control) to evaluate IFN-β and MX1 mRNA expression by RT-qPCR Individual results are shown as dots and bars represent means of at least three individual experiments. Error bars show SEM. The significance was tested by two-way ANOVA with Dunnett’s post hoc testing. **, p < 0.01, ***, p < 0.001. **C** HeLa WT or cGAS-deficient cells (a different cell line from previously, please see methods) were treated with 10 µM QVD-OPh or DMSO (solvent control) for 24, 48 or 72 h. Protein levels of MX1, p-IRF3 and p-STING were analyzed by Western blotting. GAPDH was used as a loading control. The result is representative of three individual experiments. **D** Cells from HeLa cell lines (control or STING-deficient) were treated with 10 µM QVD-OPh or DMSO (solvent control) for 72 h. Protein levels of MX1 and p-IRF3 were analyzed by Western blotting. GAPDH was used as a loading control. The result is representative of three individual experiments.

**Fig. S5.**
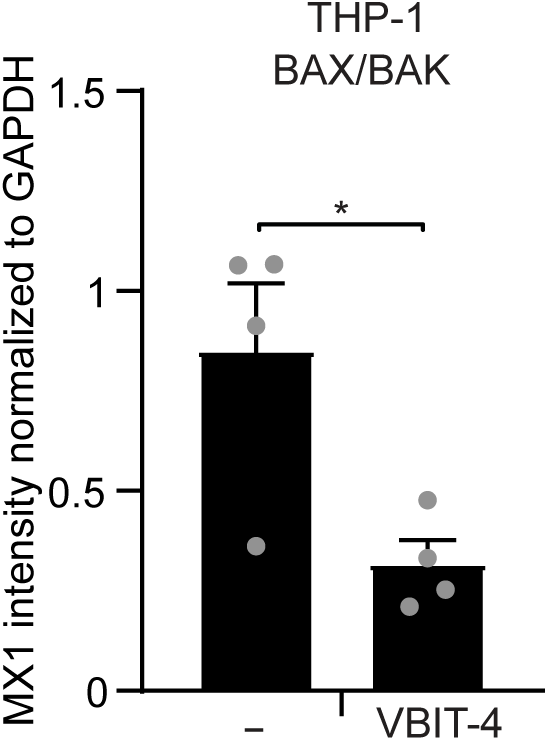
Mitochondrial DNA provides a cGAS signal in resting cells. Quantification of MX1 protein levels in THP-1 cell line (BAX/BAK CRISPR/Cas9 mutants) upon treatment with VBIT-4. MX1 intensity was measured and normalized to the loading control (GAPDH) signal using ImageJ software. Individual results are shown as dots and bars represent means of four individual experiments. Error bars show SEM. *, p < 0.05. The significance was tested by paired t-test.

**Fig. S6.**
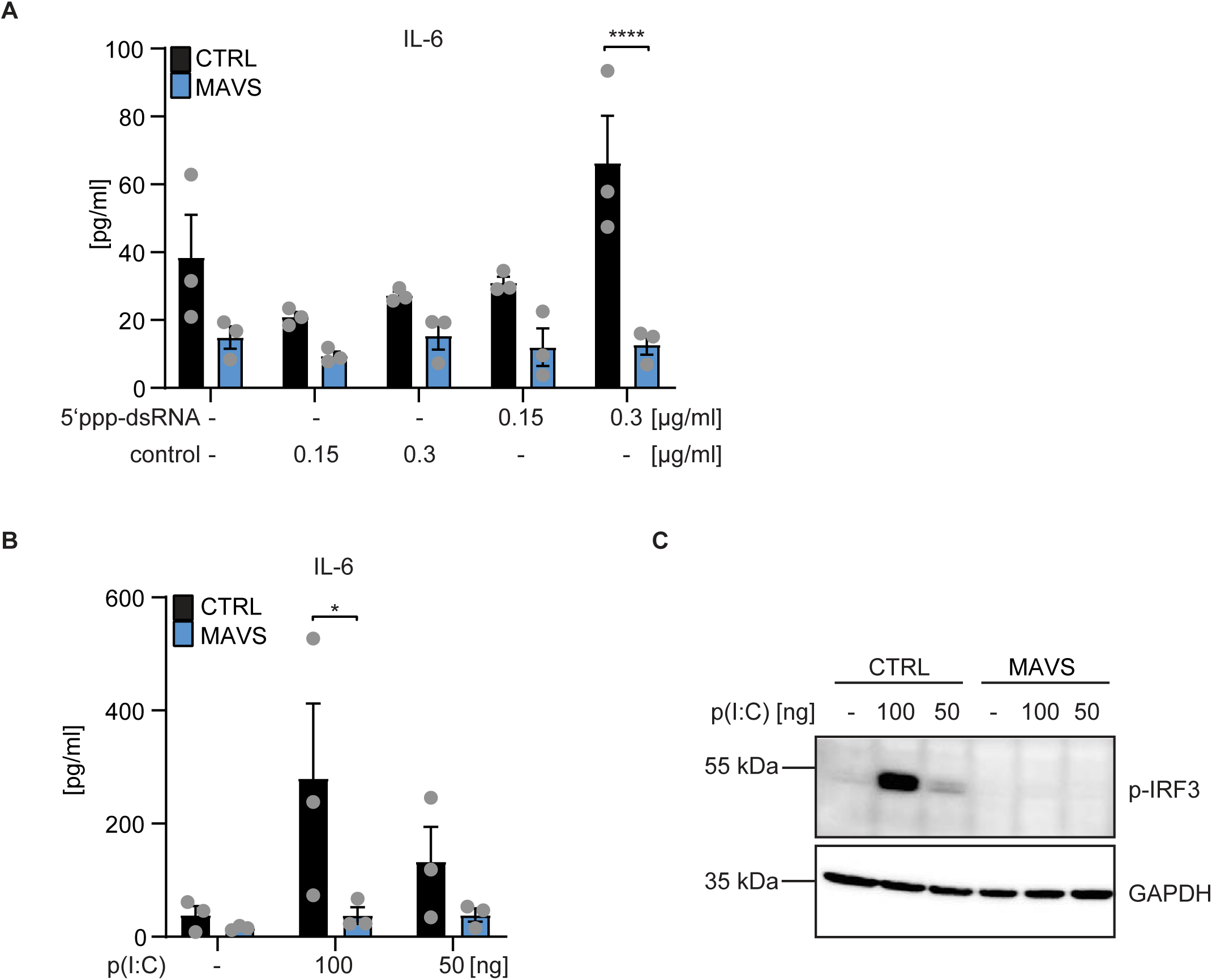
MAVS dependency of IL-6 secretion and IRF3 phosphorylation induced by RLH ligands. Cells from HeLa cell lines (control or MAVS-deficient) were transfected with 5’ppp-dsRNA (RIG-I-ligand) or control dsRNA (without 5’-triphosphate, non-stimulatory) **(A)** or transfected with 100 ng and 50 ng poly (I:C) **(B, C)**. **A, B** 24 h post-transfection, supernatants were harvested and subjected to IL-6 ELISA. Results from individual experiments are shown as symbols and bars represent means of three individual experiments. Error bars show SEM and p values were calculated by two-way ANOVA (mean comparison) statistical assay with Sidak’s multiple comparisons testing. *, p < 0.05, ****, p < 0.0001. **C** Cells were transfected with the indicated amount of poly (I:C). After 24 h, cells were lysed and levels of p-IRF3 were analyzed by Western blotting. GAPDH was used as a loading control. The result is representative of at least three individual experiments.

**Fig. S7.**
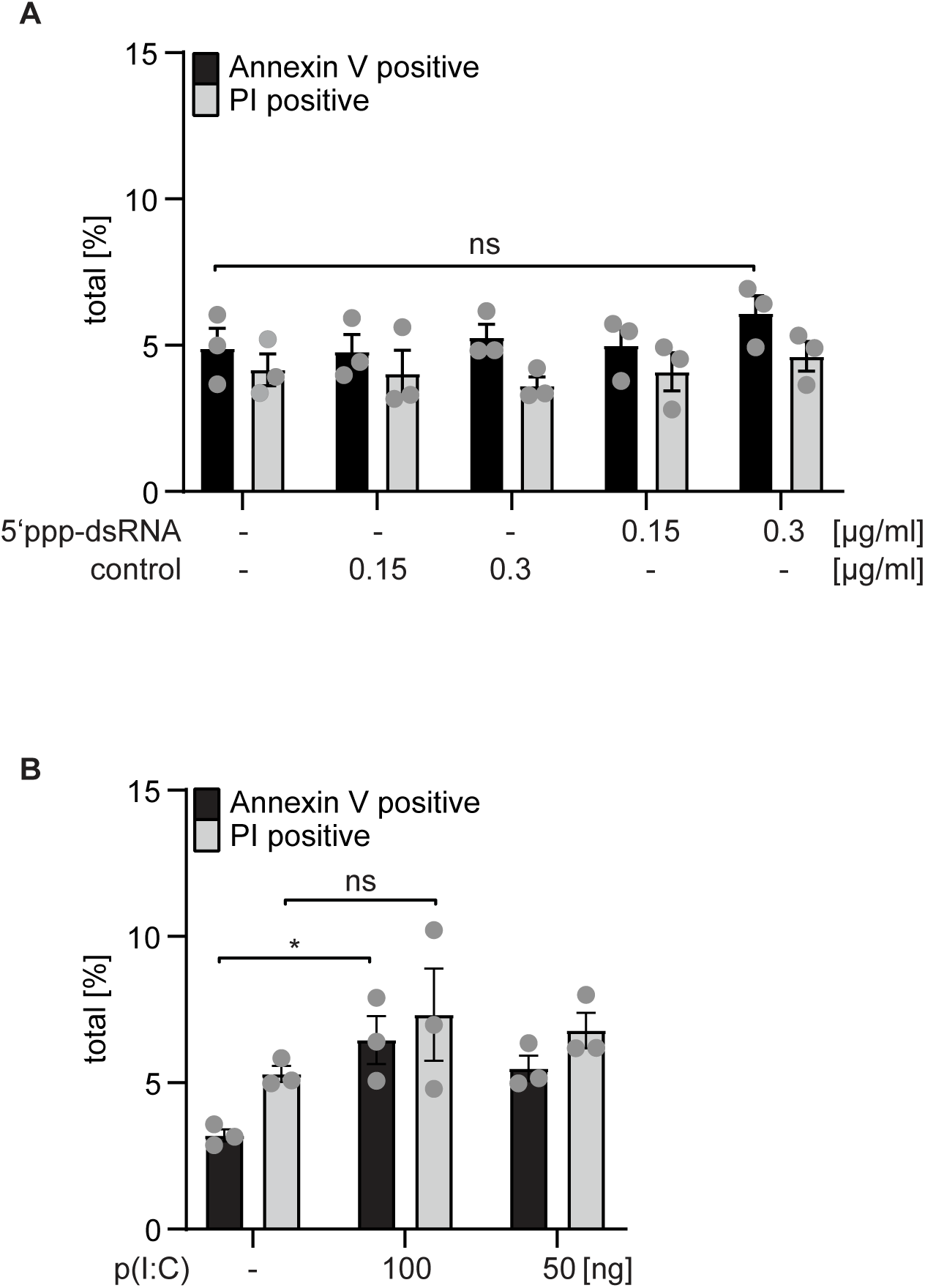
Levels of cell death following transfection of HeLa cells. Cells form HeLa cell lines (control or MAVS-deficient) were transfected with 5’ppp-dsRNA (RIG-I-ligand) or control dsRNA (without 5’-triphosphate) **(A)** or transfected with 100 ng and 50 ng poly (I:C) **(B)**. **A** 24 h post-transfection, cell death was assessed by Annexin V/PI staining. Results from individual experiments are shown as symbols and bars represent means of three individual experiments. Error bars show SEM and p values were calculated by two-way ANOVA (mean comparison) statistical assay with Dunnett’s multiple comparisons testing. Ns: p > 0.05. **B** 16 h post-transfection, cell death was assessed by Annexin V/PI staining. Results from individual experiments are shown as dots and bars represent means of three individual experiments. Error bars show SEM and p values were calculated by two-way ANOVA (mean comparison) statistical assay with Tukey’s multiple comparisons testing. *, p < 0.05.

**Fig. S8.**
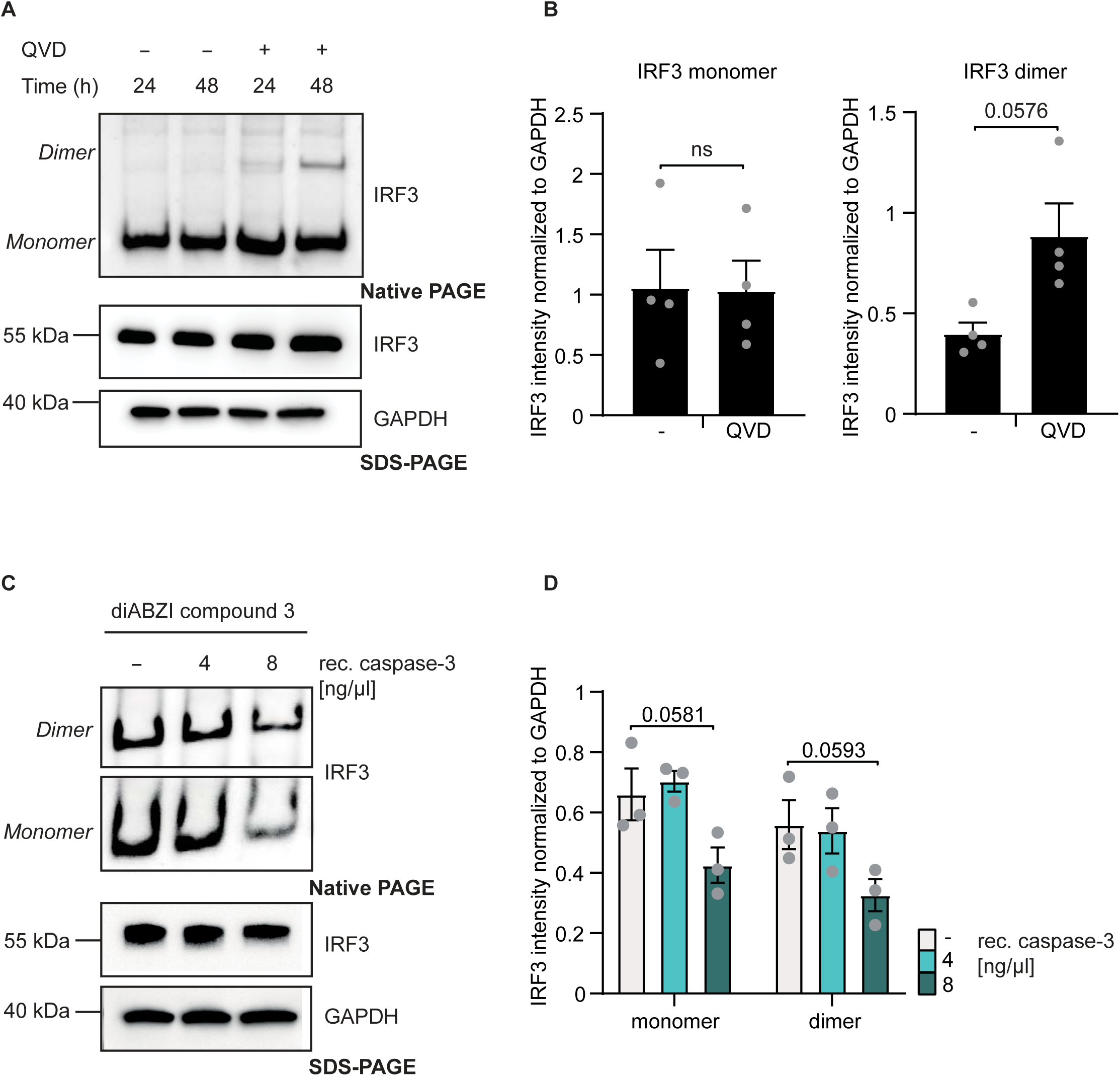
Dimerization and caspase-3 sensitivity of IRF3. **A** HeLa cells (control) were treated with 10 µM QVD-OPh or DMSO (solvent control) for 24 or 48 h. The same whole cell lysates were run on native gels to analyze IRF3 dimerization and on the SDS-PAGE to evaluate endogenous IRF3 levels. GAPDH was used as a loading control. **B** Quantification of the monomer and dimer bands of IRF3 (native gels as shown in A) 48h QVD-OPh treatment, normalized to the GAPDH (SDS-PAGE) signal. Results from individual experiments are shown as symbols and bars represent means of four individual experiments. Error bars show SEM. The significance was tested by paired t-test. Ns: p > 0.05. **C** HeLa cells (control) were treated with 2 µM diABZI compound 3 (STING agonist) for 2 h, and whole cell lysates were incubated with the indicated concentrations of recombinant caspase-3 (rec. casp-3). Samples were incubated at 37° C for 1 h. After incubation, reactions were stopped by addition of 10 µM QVD-OPh. Samples were run on the native gel to analyze IRF3 dimerization and on the SDS-PAGE to evaluate endogenous IRF3 levels. GAPDH was used as a loading control. **D** Quantification of the monomer and dimer bands of IRF3 (native gels as shown in C), normalized to GAPDH (SDS-PAGE) signal. Individual results are shown as symbols and bars represent means of three individual experiments. Error bars show SEM and p values were calculated by two-way ANOVA (mean comparison) statistical assay with Sidak’s multiple comparisons testing.

